# DNA modulates structural transitions and oligomerization kinetics of the functional amyloid CRES

**DOI:** 10.1101/2025.08.14.670404

**Authors:** Ritika Kukreja, Gail A. Cornwall, Michael P. Latham

## Abstract

Functional amyloids play critical roles in diverse physiological processes; however, the molecular mechanisms regulating their assembly remain unknown. The mouse epididymal lumen contains a functional amyloid- and nucleic acid-rich extracellular matrix with roles in host defense and sperm maturation. Cystatin-related epididymal spermatogenic (CRES), a reproductive cystatin and key component of the mammalian epididymal amyloid matrix, assembles into structurally heterogeneous amyloid forms to support the various roles of this extracellular matrix. Here, we show that CRES binds double-stranded DNA with sub-micromolar affinity in a sequence independent manner and that DNA binding accelerates amyloid formation by increasing local protein concentration and promoting specific oligomerization pathways. Using NMR spectroscopy, site-directed mutagenesis, and biophysical analyses, we find that DNA interacts primarily with the CRES loop region, thereby occluding one assembly mechanism and redirecting oligomerization through a pathway involving the L1 loop. These DNA-mediated changes in assembly kinetics and pathway selection suggest a regulatory mechanism for achieving structural and functional diversity in non-pathological amyloid systems. Our findings provide a molecular framework for understanding nucleic acid-guided amyloidogenesis and highlight how functional amyloids may exploit multiple assembly routes to fulfill their physiological roles.

## INTRODUCTION

The spontaneous aggregation of proteins can result in the formation of large supramolecular assemblies characterized by cross β-sheet (and/or cross-alpha) architectures^1^. These highly ordered structures, collectively termed amyloids, can manifest diverse morphologies including fibrillar aggregates, films, and matrices^2^. Extensive research has been conducted on pathological amyloids, predominantly associated with neurodegenerative disorders such as Alzheimer’s disease, Parkinson’s disease, Huntington’s disease, and amyotrophic lateral sclerosis^3–6^. In contrast, functional amyloids, which play crucial physiological roles, remain comparatively understudied. Non-pathological amyloids are implicated in diverse biological processes, such as cell-cell aggregation, biofilm formation, signal transduction, host defense, sperm protection and maturation, germline specification, and hormone storage^7–13^. We previously demonstrated that a functional amyloid matrix assembly with host defense functions is present in the mouse epididymal lumen^10,13,14^. This extracellular amyloid matrix is composed of several family 2 cystatins, which are a class of cysteine protease inhibitors including cystatin C, and four members of the CRES (cystatin-related epididymal spermatogenic; CRES, CRES2, CRES3, cystatin E2 in mouse) subgroup, a reproductive subgroup within the family 2 cystatins^15,16^. CRES amyloid and the endogenous epididymal amyloid matrix form different amyloid morphologies (matrix, films, fibrils) with different host defense functions (bacterial trapping vs killing) depending on the bacterial strain^13^. This study suggests that external factors can influence CRES amyloidogenesis and that the amyloid properties of CRES, and likely other CRES subgroup members, provides the epididymal amyloid matrix with functional biological plasticity.

CRES exhibits a similar mechanism of amyloidogenesis as pathological amyloids. Initially, CRES possesses a metastable globular conformation capable of rapid, reversible transitions between its native state and pre-amyloid oligomeric forms which are further influenced by cross-seeding between CRES and its various paralogs^11,17,18^. Eventually, CRES oligomerizes into large and varying matrix, film, and fibrillar structures. We have previously shown that CRES likely utilizes two distinct pathways of assembly that could account for the controlled formation of heterogeneous structures including matrices and fibril-like arrays^11,18^. One pathway of assembly involves aggregation via the L1 loop which is the critical structure involved in domain swapping for cystatin family members^19–21^, whereas the other pathway utilizes a novel mechanism unique to CRES. This latter pathway is characterized by intermolecular interactions between the flexible CRES loop (situated between β-strands 3 and 4) of one monomer and the β5 strand of an adjacent monomer resulting in the formation of a pseudo-parallel β-sheet interface between the two protomers^11^. In addition to these two pathways for amyloid assembly, external factors and changes in the physical/chemical environment of the epididymal lumen may influence CRES amyloidogenesis. Our recent studies revealed that extracellular nucleic acids are part of the epididymal amyloid matrix and contribute significantly to matrix integrity and stability^22^. Given that numerous amyloidogenic proteins, such as Aβ^23^ and SOD1^24^ amyloids, exhibit nucleic acid-templated assembly^25^, it is plausible that CRES, as a functional amyloid, may also engage in nucleic acid interactions, possibly as a mechanism to regulate the assembly and structure of the epididymal amyloid matrix.

In this present study, we elucidate the molecular interactions between CRES and DNA over a wide range of length and time scales. Employing *in vitro* fluorescence-based binding studies, we report a sub-micromolar binding affinity between CRES and double-stranded DNA (dsDNA) and investigate the stoichiometry of this interaction, revealing that multiple CRES molecules bind to small DNA constructs in a sequence independent manner. The kinetics of CRES oligomerization post-DNA interaction were characterized using dynamic light scattering (DLS), Congo red fluorescence measurements, and transmission electron microscopy and indicate that accelerated assembly of higher-order oligomers occurs in the presence of DNA. Solution-state NMR spectroscopy, complemented by site-directed mutagenesis, identified the unique CRES loop as the preferential DNA-binding interface. Subsequent oligomerization was found to proceed predominantly via interactions involving the L1 loop. NMR lifetime line-broadening experiments were utilized to analyze the assembly kinetics of early oligomeric species, and mutational studies further corroborate our hypothesis that expedited oligomerization kinetics are primarily achieved through the L1 loop. Together, these findings outline the role of DNA interactions in driving CRES amyloid assembly, offering new insights into the molecular mechanisms that underlie its amyloidogenesis and formation of a functional amyloid host defense structure.

## RESULTS

### Monomer and early oligomers of CRES bind to DNA *in vitro*

As CRES has been shown to co-localize with extracellular nucleic acids in the mouse epididymal amyloid matrix^22^, we wanted to directly investigate CRES-DNA interactions. This study used a heterologously expressed, modified version of mouse CRES (Cst 8, UniProt ID: P32766; aa 20-142 lacking the N-terminal signal sequence) that includes a cysteine to alanine mutation at residue 48 (CRES C48A; hereafter referred to as CRES) that prevents inappropriate disulfide bond formation^11^. To test if CRES binds to DNA *in vitro*, we used a fluorescence-based assay that monitors the change in polarized light emitted from a labeled DNA upon CRES binding. DNA binding was examined for three different stages of CRES amyloid assembly: monomer, “early oligomers,” and “intermediate oligomers” (Fig. 1a)^13^. Although DNA binding is not likely a two-state process, because CRES monomers are in equilibrium with early oligomers^11^, the data nonetheless fit well to a simple two-state binding model. We therefore report apparent K_d_s (K_d,app_). Monomeric and early oligomeric forms of CRES bound to a 15-basepair (bp) hairpin DNA with a K_d,app_ of 0.35 ± 0.14 μM and 0.39 ± 0.18 μM, respectively (Fig. 1b, left and center). Conversely, the intermediate oligomeric CRES showed dramatically reduced affinity for the hairpin DNA (K_d,app_ = 22.2 ± 7.3 μM) (Fig. 1b, right). Because CRES amyloidogenesis progresses over time^18^, DNA binding was monitored on the same protein samples over the course of a month. The intermediate amyloid exhibited substantial heterogeneity in its affinity for DNA over that time, whereas the binding of monomeric and early oligomeric stages of assembly changed very little (Fig. 1b). This result suggests that the monomer and early oligomers are not further progressing into intermediate oligomers under these buffer conditions over this time period. We hypothesize that the decreased DNA binding affinity for intermediate CRES oligomers may be attributed to the occlusion of DNA binding site(s) as the protein oligomerizes. A 7-bp hairpin DNA and a 40- bp dsDNA were also used as substrates in the binding assay and gave similar results as observed for the 15-bp DNA hairpin: monomeric CRES displayed a K_d,app_ = 0.54 ± 0.08 μM and 0.74 ± 0.23 µM and early oligomers exhibited a K_d,app_ = 0.32 ± 0.02 μM and 0.82 ± 0.25 µM for 7-bp hairpin and 40-bp dsDNA substrates, respectively (Supplementary Fig. 1). For both DNA substrates, DNA binding to intermediate oligomers was also decreased. These binding reactions followed the same month-long trends as observed with the 15-bp hairpin DNA (Supplementary Fig. 1a). Lastly, CRES binding to DNA appears to not be sequence specific.

**Figure 1.**
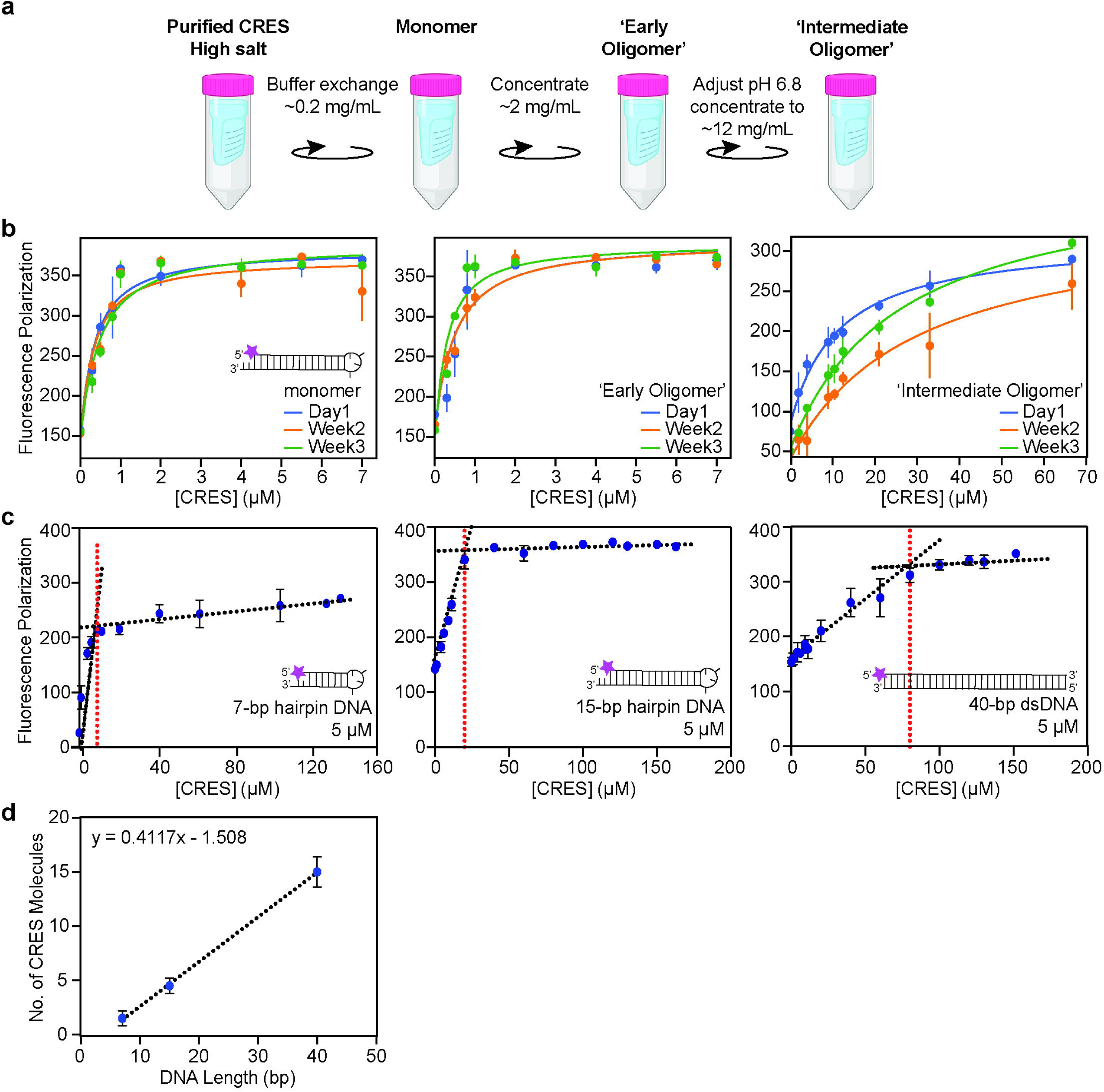
Multiple CRES molecules bind to DNA with nanomolar affinity. (a) Preparation of CRES amyloidogenesis intermediates. (b) Change in polarization of a fluorescently labeled 15-bp hairpin DNA as a function of the concentration of CRES for different stages of assembly. The solid lines represent the fitting of the data to a two-state binding isotherm. (c) Fluorescence polarization-based stoichiometric analysis for CRES binding to dsDNA substrates. Dashed black lines represent linear fits to the data, and dashed red line denotes the saturation of binding. (d) Correlation between number of CRES molecules binding to the DNA length from (c). All data points are displayed as the mean and standard deviation of three separate experiments.

The fluorescence polarization binding assay was then used to investigate the stoichiometry of the interaction between CRES and DNA. The DNA saturation curves, determined with a constant DNA concentration of 10 – 20x the K_d,app_ in an active site titration experiment, indicated that ∼1-2 CRES molecules bind to a single 7-bp hairpin DNA, ∼4-5 CRES molecules bind to a single 15-bp hairpin DNA, and ∼14-16 CRES molecules bind to the 40-bp dsDNA (Fig. 1c). Thus, as the length of the DNA increases, there is a corresponding increase in the number of CRES molecules that bind to the DNA. The slope of the number of CRES molecules bound versus the number of base pairs of dsDNA (Fig. 1d) suggests that each CRES binds to about 3-4 base pairs of dsDNA. Together, the DNA binding data support a hypothesis that CRES molecules bind to independent binding sites along dsDNA, perhaps to act as an initial template for CRES amyloidogenesis.

### NMR reveals DNA-driven conformational exchange and complex formation by CRES

To understand the specific interactions between CRES and DNA, we used solution state NMR, since the monomeric protein (14.6 kDa) is soluble^11^. Two types of perturbations were observed in 2D ^15^N,^1^H TROSY-HSQC correlation spectra upon the titration of the 15-bp hairpin DNA into a ^15^N-labeled CRES sample (Fig. 2 and Supplementary Fig. 2): line broadening (i.e., loss of signal, Fig. 2a) and changes in the chemical shift position and intensity of peaks upon DNA binding (Fig. 2b). Line broadening could occur if interactions with DNA caused the formation of larger CRES_n_-DNA complexes that tumble more slowly in solution; alternatively, line broadening could be the result of DNA binding inducing µs-ms timescale conformational exchange within specific regions of CRES, as previously described^11^. Residues that experienced moderate line broadening predominantly mapped to the L1 loop and proximal beta stands (e.g., Q77, I78, T79, D80, R81, E83) and the α-helix (e.g., V45, A43) (Fig. 2c) – regions of the protein suspected to be involved in a domain-swapping assembly pathway. On the other hand, chemical shift perturbations (CSPs) coupled with changes in peak intensity are consistent with an intermediate exchange phenomenon. For the residues undergoing intermediate exchange, DNA titration caused their peak positions to change and their intensities to decrease until 20-30% M equivalents (M eq) of DNA was added to the protein. At this point these peaks lost all intensity. Upon addition of >30% DNA, these CRES peaks re-appeared at new chemical shift positions but with less intensity compared to CRES without DNA, again consistent with the formation of larger CRES_n_-DNA complexes. Residues exhibiting intermediate exchange mapped to the CRES loop (L99, N100, N101, T102, N104, C105, K109) and β4- and β5-strand regions (E131) (Fig. 2c). Because the CRES loop shows intermediate exchange CSPs, the NMR titration results suggest that CRES largely interacts with DNA through the CRES loop.

**Figure 2.**
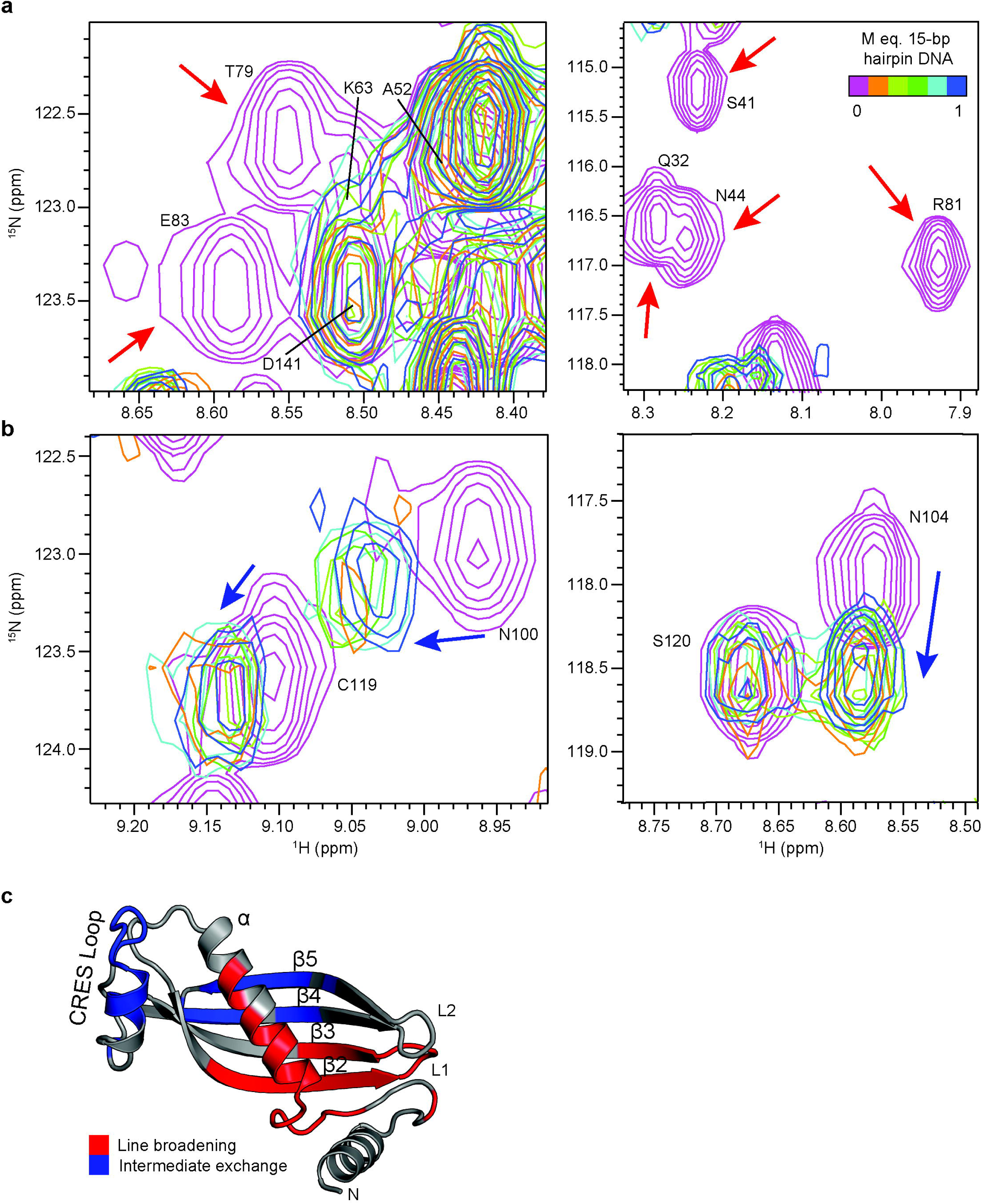
Solution-state NMR analysis of CRES binding to DNA. Overlay from regions of the 2D ^15^N,^1^H TROSY-HSQC spectra of ^15^N-labeled CRES with increasing concentrations of DNA. The overlaid spectra show resonances undergoing extreme line-broadening (a; highlighted by red arrows), and resonances undergoing intermediate exchange (b; highlighted by blue arrows). (c) Perturbations mapped onto the monomeric structure of CRES (PDB ID: 6UIO)^11^ illustrating residues that experience line-broadening (red) or intermediate exchange (blue).

Next, we compared the titration of ^15^N-labeled CRES with a 7-bp hairpin DNA to the titration with the 15-bp hairpin DNA (Supplementary Fig. 3). Similar CSPs were observed as with the longer hairpin; however, residues that previously showed large decreases in peak intensities at 20- 30% M eq of 15-bp hairpin were now less line-broadened. With the 7-bp hairpin DNA, extreme line broadening did not occur until 40-50% M eq of DNA was added (Supplementary Fig. 3a). Because of the better signal-to-noise arising from forming smaller complexes, we observed an additional type of perturbation: fast exchange behavior (i.e., the population-weighted average peak position) with decreasing peak intensity as the CRES_n_-DNA complexes become more populated (Supplementary Fig. 3d). These peaks correspond to residues in the β3-strand and L1 and L2 loops.

In these two NMR titration experiments, we observe monomeric CRES exchanging with larger CRES_n_-DNA complexes. This exchange likely involves a heterogeneous mixture of complexes with varying stoichiometries: n = 1, 2, 3, 4, or 5 CRES for the 15-bp hairpin, and n = 1 or 2 CRES for the 7-bp hairpin. At ∼20-30% M eq of 15-bp hairpin DNA or 40-50% M eq of 7-bp hairpin DNA, nearly all the available monomeric CRES is bound to DNA, as confirmed by our stoichiometric analysis (Fig. 1c), where the stoichiometry saturation threshold coincides with the observed line broadening in the NMR experiments. The observed line broadening results from ¹LJN lifetime effects; as the biomolecular complexes increase in size, they tumble more slowly in solution, leading to faster relaxation and consequently broader linewidths. The extent of line broadening therefore reflects the size of the largest complex formed upon DNA saturation. Thus, saturation of the 15-bp hairpin DNA produces more pronounced line broadening in the monomeric CRES spectrum, indicative of larger CRES_5_-DNA complexes, compared to the less extensive broadening observed with 7-bp hairpin DNA saturation, which corresponds to smaller CRES_2_-DNA complexes.

### DNA binds to the CRES loop

To obtain molecular models of DNA-bound CRES, we employed the rigid-body docking program HADDOCK^26,27^. The residues in intermediate exchange in the DNA titration spectra were defined as “active” residues in HADDOCK for docking the 15-bp hairpin DNA with monomeric CRES. As expected, most of the resulting HADDOCK models showed interactions between the CRES loop and the major or minor groove of the hairpin DNA (Supplementary Fig. 4a). Additionally, the 15-bp hairpin DNA was docked with a L1 loop-mediated domain-swapped dimer model of CRES, also created with HADDOCK (see methods), again using the CRES loop and β5-strand region as active residues. These domain-swapped dimer models not only showed protein-DNA interactions in the CRES loop but displayed additional interactions between the DNA and residues in the antiparallel β-strands that form upon domain-swapping (e.g., R81 and K75) (Supplementary Fig. 4b).

Next, a series of negatively charged CRES mutants, designed based on the CRES-DNA interactions in the HADDOCK models, were assessed for their effect on DNA binding using the fluorescence polarization binding assay (Fig. 3). Specifically, groups of polar residues in the CRES loop were mutated to either aspartate (N94D/N101D/T102D/N104D; “loop mutant D” or LMD) or glutamate (K109E/K110E) (Fig. 3a). Two β2-strand charge reversal mutants (K75E and R81E) were also generated to disrupt potential DNA-CRES interactions identified in the domain-swapped HADDOCK models (Fig. 3b). Decreased binding affinity for the 15-bp hairpin DNA was observed with LMD (K_d,app_ = 4.2 ± 1.3 μM; 12x greater than wild type) and K109E/K110E (K_d,app_ = 0.58 ± 0.13 μM; 1.7x greater than wild type) mutants in the monomeric form of CRES (Fig. 3c). These results are consistent with polar and positively charged residues in the CRES loop interacting with the negatively charged phosphate backbone of the DNA. Decreased binding affinities were also observed for monomeric CRES with the K75E (K_d,app_ = 1.5 ± 0.36 μM; 4.3x greater than wild type) and R81E (K_d,app_ = 0.82 ± 0.06 μM; 2.3x greater than wild type) mutations that are adjacent to the L1 loop involved in the domain-swapping pathway of assembly (Fig. 3c). In the domain-swapped dimer HADDOCK models, K75 and R81 become part of the new antiparallel β-sheet (Fig. 3b and Supplementary Fig. 4b), and their interactions with dsDNA suggest a possible role for DNA binding in either facilitating the formation of and/or stabilizing the domain-swapped dimer.

**Figure 3.**
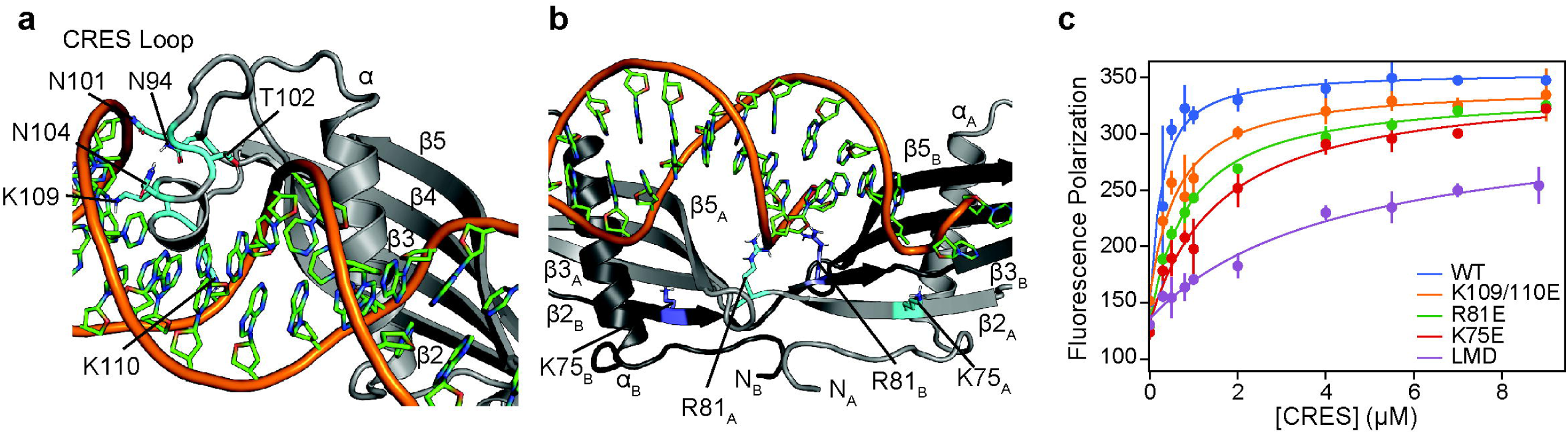
CRES primarily binds DNA through the CRES loop. (a) HADDOCK model of CRES monomer bound to DNA highlighting interactions between residues in the CRES loop (N94, N101, T102, N104, K109, and K110) and the backbone of the DNA (orange). (b) HADDOCK model of domain-swapped CRES dimer bound to DNA highlighting interactions between K75 and R81 and the DNA. (c) Polarization of a fluorescently labeled 15-bp hairpin DNA as a function of the concentration of CRES mutants. The data points are displayed as the mean and standard deviation of three replicates, and the solid lines represent the fitting of the data to a two-state binding isotherm.

### DNA facilitates CRES oligomerization

To understand the effect of CRES-DNA binding on oligomerization, we employed dynamic light scattering (DLS) to measure the change in particle size for CRES molecules in the absence and presence of a 1:1 molar ratio of 15-bp hairpin DNA. In the absence of DNA, CRES monomers had an initial diameter of 4.62 ± 0.48 nm at day 0, which was unchanged by day 14 (4.9 ± 0.5 nm) (Fig. 4a). CRES initially has a population of aggregates ∼220 nm in diameter, and larger aggregates are observed by day 8 (diameter ∼482 ± 32 nm) and day 14 (diameter ∼1285 ± 151 nm) (Fig. 4a; Supplementary 5), consistent with our previous results^13,18^. Conversely, CRES molecules incubated with equimolar concentrations of 15-bp hairpin DNA initially exhibited a larger monomeric state (7.18 ± 0.65 nm) and similar aggregates (188 ± 16 nm) at day 0. This monomeric population was lost at day 4, and there was an accelerated transition to much larger oligomeric species (1718 ± 176 nm at day 8 and 4801 ± 835 nm) at day 14 (Fig. 4a). These findings confirm our hypothesis that DNA interactions facilitate rapid protein assembly leading to oligomerization.

**Figure 4:**
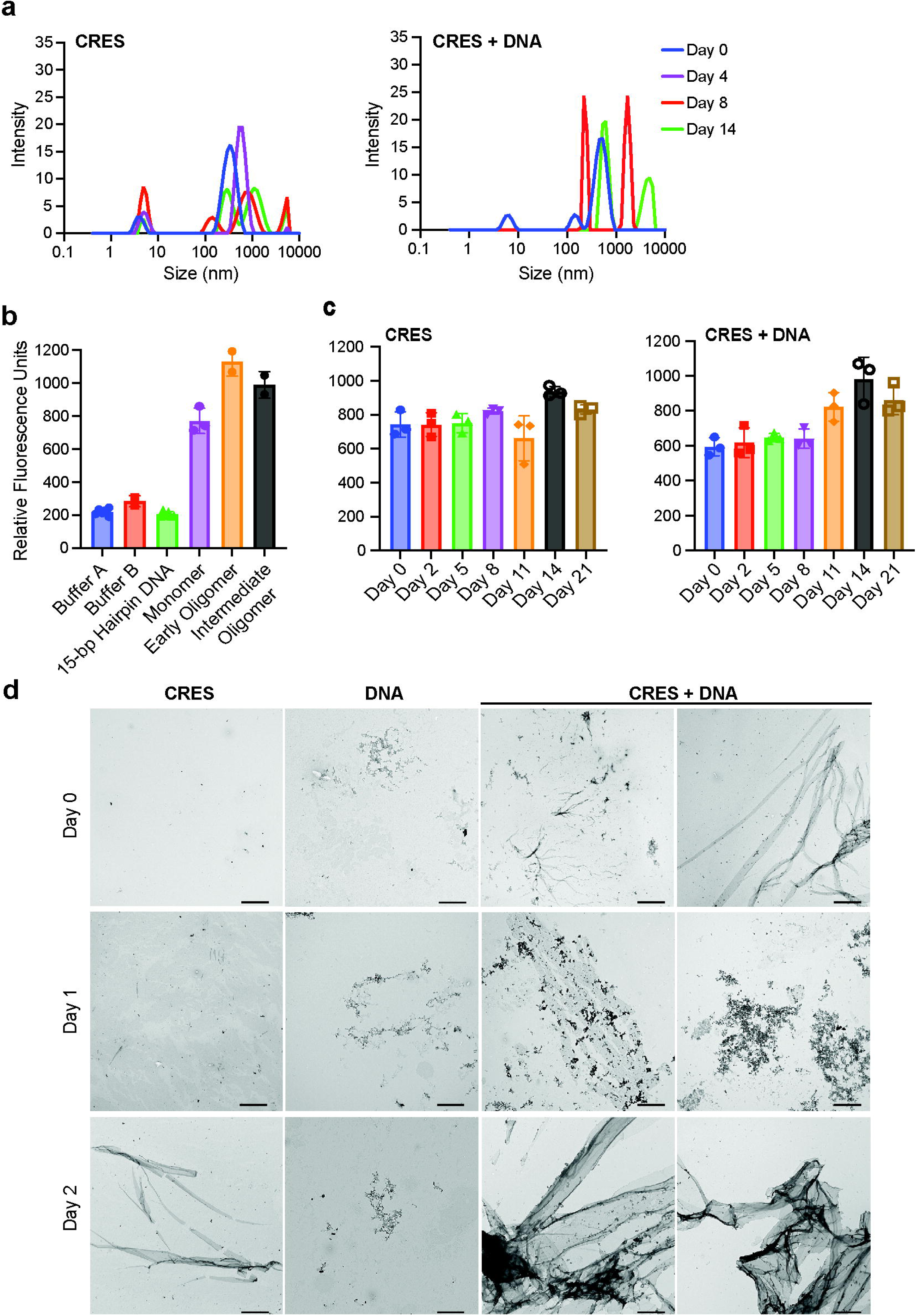
CRES binding to DNA facilitates oligomerization. (a) Dynamic light scattering size distribution by intensity for monomeric CRES samples without (left) and with (right) equimolar 15-bp hairpin DNA. (b) Congo red fluorescence controls for monomer/’early oligomer’ buffer (Buffer A: 25 mM MES pH 6), ‘intermediate oligomer’ buffer (Buffer B: 25 mM MES, 50 mM HEPES pH 6.8), and 15-bp hairpin DNA only as well as for different stages of CRES assembly. (c) Congo red fluorescence measurements of monomeric CRES samples in the absence (left) and presence (right) of equimolar concentrations of 15-bp hairpin DNA. (d) Negative stain TEM of CRES and 15-bp hairpin DNA alone and after 60 min (Day 0), 24 hrs (Day 1), and 48 hrs (Day 2) co-incubation at room temperature. Scale bar, 2 µm.

To further validate amyloid formation and protein aggregation resulting from CRES-DNA interactions, we conducted fluorescence-based assays to assess the binding capacity of CRES oligomers to Congo red, an amyloid-specific dye, in the absence and presence of DNA (Figs. 4b and 4c). Conventional ThT fluorescence measurements were initially attempted; however, increased fluorescence was observed when the dye was incubated with DNA, indicating that ThT was binding to the DNA rather than specifically to the protein aggregates, due to the positively charged nature of ThT. Control experiments confirm an increase in Congo red fluorescence as CRES progresses from monomer to intermediate oligomers (Fig. 4b) verifying that this dye is sensitive to the early stages of CRES amyloidogenesis. Initially, monomeric CRES incubated with DNA showed lower fluorescence intensity (∼640 relative fluorescence units - RFUs) compared to CRES alone (∼740 RFUs) which is likely due to electrostatic repulsion between the negatively charged phosphate backbone of the DNA and the anionic Congo red dye (Fig. 4c). But, as time progressed and larger oligomers formed, the fluorescence intensity increased for CRES samples incubated with DNA (∼860 RFUs) (Fig. 4c), whereas the fluorescence intensity for apo CRES remained essentially the same (∼830 RFUs). Quantitatively, the change in relative fluorescence over a 21-day incubation period was substantially greater for samples containing equimolar concentrations of DNA compared to samples without DNA (ΔRFU∼ 220 vs 90, respectively; Fig. 4c), indicating that DNA accelerates the formation of larger CRES oligomers. Thus, the outcomes observed in DLS and Congo red experiments were consistent with each other and support our hypothesis that DNA interactions promote the formation of larger CRES aggregates.

To visualize the macroscopic structural changes occurring to the protein upon interactions with DNA, we used negative-stain transmission electron microscopy (TEM) imaging. The incubation of CRES monomers with 15-bp hairpin DNA resulted in the rapid formation of large assemblies including thin film fragments at 0 days (60 min incubation) that progressed to matrix arrays and elaborate films after 1 and 2 days, respectively. In contrast, after 2 days CRES in the absence of DNA remained as a thin granular layer with only small fragments of film (Fig 4d). Together with the stoichiometry experiments (Fig. 1d), these data demonstrate that DNA binding facilitates amyloidogenesis by increasing primary CRES nucleation, likely through the local concentration of CRES monomers that promotes the rapid growth of larger oligomers.

### The CRES monomer exchanges with large, oligomeric DNA-CRES complexes

The kinetic processes associated with DNA binding and subsequent CRES assembly were studied with ^15^N spin relaxation NMR experiments. These experiments generally monitor molecular motions on the picosecond to nanosecond timescales and sometimes slower motions (microsecond to millisecond) through exchange processes^28,29^. To observe the possibility of monomeric CRES exchanging (i.e., coming on and off) with large CRES_n_-DNA complexes or even larger oligomeric species (e.g., [CRES_n_-DNA]_x_ or [CRES_n_-DNA]+xCRES), we used small DNA:CRES ratios of 2.5% and 5% M eq of 15-bp hairpin DNA. Under these conditions, the concentrations of the large complexes are much lower than that of the monomer and therefore line broadening is minimized. For wild-type CRES, the addition of 15-bp hairpin DNA caused an increase in the ^15^N transverse relaxation rate (R_2_) in a concentration-dependent manner as the average ^15^N R_2_ increased from 20.5 s^-1^ for CRES alone to 32.1 s^-1^ and 36.7 s^-1^ after the addition of 2.5% and 5% M eq of DNA, respectively (Fig. 5a). This increase in R_2_ is much larger than what is expected from the addition of sub-stoichiometric amounts of DNA. For a 1:1 CRES-DNA complex, the ^15^N R_2_ rates would be ∼20-25 s^-1^ based on a CRES-DNA molecular weight of ∼26 kDa^30^. Instead, the ^15^N R_2_ rates of 32.1 s^-1^ and 36.7 s^-1^ support the assertion that monomeric CRES is exchanging with a much larger complex (i.e., CRES_n_-DNA; Fig 5b).

**Figure 5.**
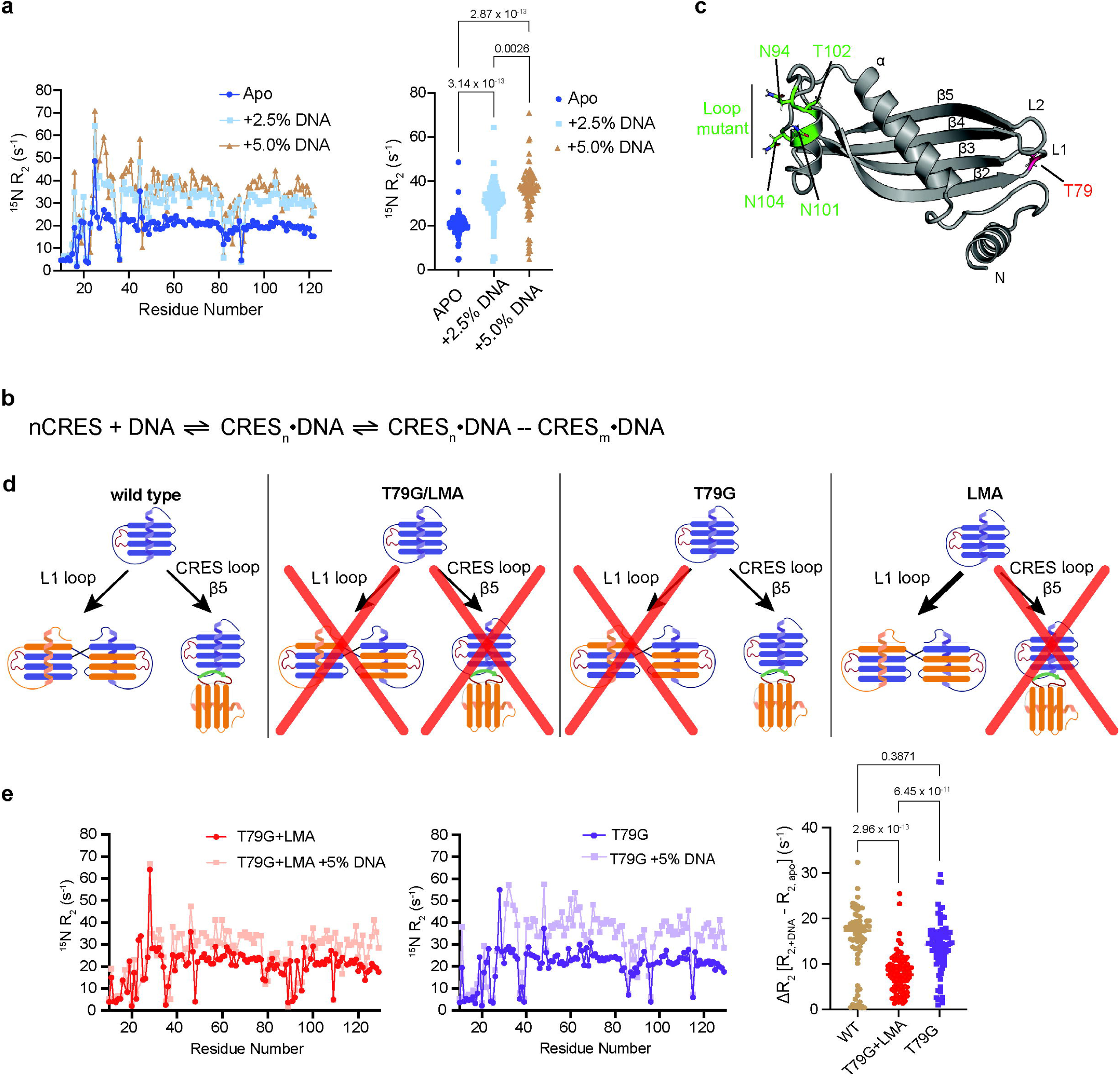
CRES exchanges with larger DNA-bound CRES oligomers. (a) Comparison of backbone amide ^15^N transverse relaxation rates (^15^N R_2_) of wild-type CRES in the absence and presence of 2.5% and 5% hairpin DNA. P-values were calculated by one-way ANOVA and Tukey’s multiple comparisons test. (b) Schematic of the proposed DNA-CRES binding mechanism. (c) Model (PDB ID: 6UIO) highlighting the mutants used to disrupt the two oligomerization pathways. (d) Cartoon depicting the pathway blocking mutants: T79G (blocks L1 loop pathway), LMA (blocks CRES loop interactions), and T79G+LMA (blocks both pathways). (e) Comparison of ^15^N lifetime line-broadening (^15^N ΔR_2_) measurements for wild type, T79G+LMA, and T79G CRES in the presence of 5% 15-bp hairpin DNA. P-values were calculated by one-way ANOVA followed by Tukey’s multiple comparisons post-test.

To understand the exchange dynamics between CRES and DNA-bound CRES, ^15^N lifetime line-broadening^28,31^ (^15^N ΔR_2_) was next measured at these low DNA:CRES ratios. ^15^N ΔR_2_ arises from the association and dissociation of the fast-tumbling monomeric CRES, which has smaller R_2_ rates and is observable in the NMR spectra, and the slow-tumbling CRES_n_-DNA or larger oligomeric species, which have much larger ^15^N R_2_ rates and are therefore invisible in the NMR spectra. ^15^N ΔR_2_ was determined from the difference in ^15^N R_2_ in the presence and absence of exchange (i.e., in the presence and absence of sub-stoichiometric DNA). The kinetics (k_ex_) and populations for the exchange process of monomeric CRES interacting with DNA-bound species magnetic field strengths (i.e., 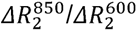 ratio at 850 MHz and 600 MHz ^1^H frequency)^28,32^. (CRES_n_-DNA) can be calculated from the ratio of ^15^N ΔR_2_ values collected at two static The interaction of CRES with the CRES_n_-DNA complex was found to be in fast exchange (α = 0.95) on the timescale of transverse relaxation (i.e., k_ex_ >> R_2,bound_). Because the exchange process is in fast exchange, the bound state population (p_B_) can be approximated through ΔR_2_ ≍ p_B_R_2,bound_ with an estimate for R_2,bound_, which is the transverse relaxation rate for the slowly tumbling large oligomeric species. In a two-state model (i.e., 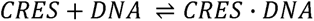), R_2,bound_ for the 1:1 CRES·DNA complex (∼26.6 kDa) would be ∼20 – 25 s^-1^, resulting in a p_B_ ∼0.45 and ∼0.65 for the 2.5% and 5% DNA added samples. However, the expected p_B_ is 0.025 or 0.05 based on the 2.5% or 5% concentration of DNA added to the sample for a 1:1 complex. Therefore, this NMR-derived population is inconsistent with a simple two-state binding model and further supports the hypothesis that the presence of DNA results in large, oligomeric species.

If monomeric CRES is exchanging with a CRES_5_-DNA complex (∼84 kDa; Fig. 5b) with an estimated 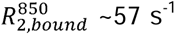, the population of DNA-bound CRES molecules would be ∼0.21 and ∼0.30 for the 2.5% and 5% addition of DNA, respectively. These p_B_s suggests that 6-8 CRES molecules interact with the 15-bp hairpin DNA (i.e., 0.21/0.025 ≈ 8 and 0.3/0.05 ≈ 6), which is closer but still exceeds the expected of 4-5 CRES per DNA based on the fluorescence stoichiometry experiment (Fig. 1c). Again, this discrepancy indicates that NMR is detecting exchange with even larger complexes in addition to the initial CRES_n_-DNA complex. Nevertheless, using this p_B_ and k_ex_ for the addition of 5% DNA to the sample (0.3 and 840 s^-1^, respectively), the k_on_ and k_off_ rate constants were estimated to be 2.7 x 10^9^ s^-1^ M^-1^ and 5.5 x 10^2^ s^-1^, respectively. Consistent with the effects of the charge reversal mutants (Fig. 3), we note that the on-rate is faster than diffusion and underscores a role for electrostatic attraction between CRES and the DNA, as is seen for other protein-nucleic acid complexes^33^. These results indicate that the activation energy associated with CRES-CRES complex formation is decreased in the presence of DNA which further corroborates our hypothesis that interactions with DNA facilitate the protein aggregation process.

### Blocking both CRES assembly pathways limits oligomerization in the presence of DNA

To determine the nature of the large, slowly relaxing CRES_n_-DNA complex, we designed a mutant that would simultaneously obstruct both the L1 and CRES loop assembly pathways while maintaining a DNA binding affinity similar to wild type (Fig. 5c). Domain-swapping in many cystatins results in the formation of a long intermolecular β-strand between two monomers and occurs through a conserved motif in the L1 loop between the β2 and β3 strands (Q-X-V-X-G, where X is any residue; Fig. 5d)^34,35^. Within this motif, the valine is the most critical residue: the preferred Ramachandran angles for valine do not support the hairpin turn between the β2-β3 strands thereby leading to domain swapping^35,36^. Crystallographic studies on cystatins show that mutating this conserved residue in the L1 loop to a glycine or asparagine prevents domain swapping, since the phi/psi angles for glycine and asparagine allow the hairpin turn orientation^35,36^. CRES has a modified version of this motif in its L1 loop (Q-I-T-D-R) with the threonine residue (position 79) at the apex of the loop instead of valine. The phi/psi angles for this threonine occupy a strained, but not completely unfavorable, conformation in Ramachandran space which may stabilize the monomeric form of CRES and reduce the probability of domain swapping as compared to other family 2 cystatins such as cystatin C^11^. Therefore, we mutated threonine 79 of the CRES L1 loop to a glycine (T79G) to mimic the mutants that facilitate monomeric cystatin conformations. The monomeric CRES T79G mutant showed similar binding affinity (K_d,app_ = 0.25 ± 0.02 μM) for the 15-bp hairpin DNA as wild type (Supplementary Fig. 6). Previous studies have shown that the CRES loop mutant N94A/N101A/T102A/N104A (LMA), designed to block the CRES loop interactions involved in oligomerization, exhibited different structural assemblies over a shorter time period compared to wild type CRES, which suggests a role for the CRES loop in CRES oligomerization (Fig. 5d)^11^. This LMA mutant also bound the 15-bp hairpin DNA with similar affinity (K_d,app_ = 0.27 ± 0.04 μM; Supplementary Fig. 6) as wild type CRES.

Since LMA and T79G both bind to DNA with wild type affinity but alter the CRES loop and L1 loop oligomerization pathways, respectively, we generated the T79G+LMA mutant (T79G/N94A/N101A/T102A/N104A; Figs. 5c and 5d) and examined its exchange properties in the NMR in the presence of sub-stoichiometric DNA. In the presence of 5% DNA, the T79G+LMA mutant had an average ΔR_2_ of 3.9 s^-1^ which is considerably less than the average ΔR_2_ = 14.4 s^-1^ observed for wild type CRES with 5% DNA (Fig. 5e). Using the relation ΔR_2_ ≍ p_B_R_2,bound_ and an R_2,bound_ ∼57 s^-1^ for the 84 kDa CRES_5_-DNA complex, a p_B_ ∼0.15 is calculated. If 5 CRES protomers bound to a single DNA, then the population of DNA-bound CRES would be 0.2 with 5% DNA in the sample which is consistent with the p_B_ ∼0.15 from the NMR results. Thus, the T66G+LMA mutant successfully blocks both CRES assembly pathways and results in a ‘stable’ CRES_5_-DNA complex – that is additional CRES molecules are not binding to the complex via CRES-CRES interactions.

To test the contributions of each oligomerization pathway on DNA-facilitated oligomerization, we next measured ^15^N spin relaxation on the T79G (blocked L1 loop) and LMA (blocked CRES loop) mutants separately (Fig. 5d). For T79G CRES, an average ΔR_2_ of 12.2 s^-1^ at 600 MHz was obtained (Fig. 5e), which is comparable to the value obtained for wild type (14.4 s^-1^ at 600 MHz) and yields a k_ex_ ∼223 s^-1^. On the other hand, although ^15^N R_2_ could be measured on CRES LMA alone, extreme line-broadening was observed after the addition of 5% 15-bp hairpin DNA to the ^15^N-labeled LMA sample, likely because CRES quickly proceeded to very large oligomeric species (Supplementary Fig. 7). This observation is consistent with our previously published DLS results showing that the CRES LMA mutant does not form a stable oligomer and rapidly assembles into large aggregates^11^.

Finally, as a last piece of evidence for the role of DNA-induced L1-mediated CRES oligomerization, the peak intensities of HSQC spectra were monitored over time. The formation of later stages of large CRES assemblies (i.e., post formation of CRES_n_-DNA complex over a time period of “days”) should lead to species too large to be observed in HSQC spectra. In the absence of DNA, wild type CRES lost ∼20% of its intensity after 24 days (Fig. 6a, Supplementary Fig. 8a). However, in the presence of 5% DNA, wild type CRES lost ∼50% of its intensity after only ∼12 hours and continued to lose intensity at a slower rate over the next ten days (Fig. 6a, Supplementary Fig. 8b). When the L1 loop pathway was blocked with the T79G mutant, the intensity loss in the presence of 5% DNA was the same as wild type CRES without DNA at seven days, and then the intensity loss accelerated to ∼50% of the signal at 20 days (Fig. 6a, Supplementary Fig. 8c). The slower rate of peak intensity loss and thus oligomerization observed in the presence of DNA in this L1 loop pathway-blocking mutant demonstrates the role of this pathway in advanced oligomer formation. These data confirm that the later stages of advanced CRES oligomerization proceed through L1-mediated oligomerization and that DNA binding greatly accelerates that overall process.

**Figure 6.**
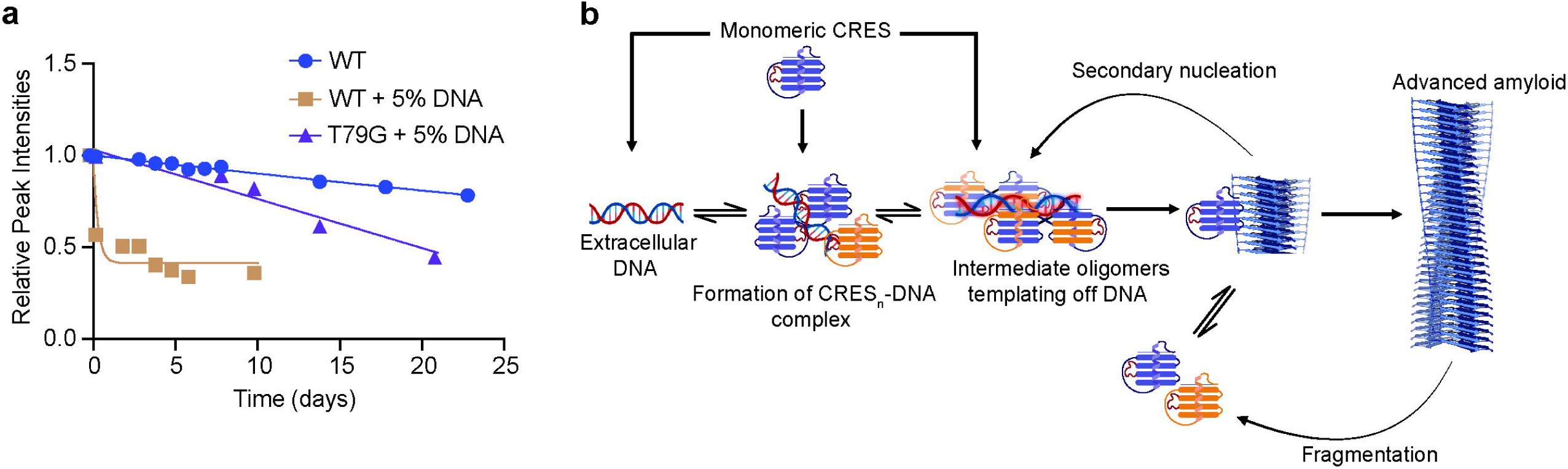
DNA facilitates the rapid formation of large CRES aggregates. (a) Global peak intensity loss relative to wild type is plotted as a function of time (in days) to compare rate of oligomerization between wild type, wild type +5% DNA, and T79G mutant +5% DNA CRES ^15^N NMR samples. (b) Proposed mechanism of DNA-influenced CRES amyloidogenesis. CRES preferentially binds the nucleic acid phosphate backbone through its unique CRES loop, stabilizing the L1 loop-mediated domain swapped dimer conformation. Increased proximity and local concentration of CRES molecules upon DNA binding promote rapid oligomerization. Once critical nucleus formation occurs, rapid assembly of β-sheet-rich amyloid fibrils proceeds (elongation phase). Secondary mechanisms including surface-mediated nucleation on pre-formed fibrils or fibril fragmentation may serve as additional determinants of the fate of the assembly.

Overall, the T79G+LMA, T79G, and LMA mutants provide insights into the interplay between different CRES assembly pathways and the effects that DNA have on each (Fig. 6b). Our binding data demonstrate that multiple CRES protomers bind to DNA independently via the CRES loop. When both assembly pathways are blocked (T79G+LMA mutant) only early CRES_n_-DNA complexes were formed without further higher-order assembly, as evidenced by low ^15^N ΔR_2,obs_ (3.9 s^-1^) for the T79G+LMA mutant. When only the CRES loop pathway was blocked, CRES-DNA complexes bypass the early metastable oligomer and rapidly formed large aggregates by the L1 loop/domain swapping pathway, as evident by the extreme NMR line-broadening and signal loss for LMA CRES in the presence of 5% DNA. When the cystatin-like L1 loop pathway was blocked (T79G mutant), intermediate oligomers can form, as evident by ΔR_2,T79G_ ≅ ΔR_2,WT_, but the formation of larger assemblies is prevented, as shown by the slower loss of NMR peak intensities after 20 days. These results support our hypothesis that upon DNA binding, the CRES loop facilitates intermediate oligomer formation, followed by L1 loop-mediated aggregation into larger oligomeric species.

## DISCUSSION

The study of amyloid-nucleic acid interactions has expanded across multiple systems, including disease-associated amyloids like amyloid-beta (Aβ)^23,37,38^ and α-synuclein^39,40^, as well as functional bacterial amyloids such as Hfq and RepA^41–45^. Collectively, these results illuminate conserved mechanisms in nucleic acid-facilitated amyloid formation and suggest a common molecular framework that underlies these interactions. Here, we provide significant insights over a wide range of time and length scales into the molecular mechanisms underlying the interactions between CRES, a functional amyloid protein, and DNA. Our findings reveal a sophisticated interplay between DNA binding, multiple assembly pathways, and amyloidogenesis. Our data also show similar characteristics between CRES-DNA and other amyloid-nucleic acid complexes such as electrostatic complementarity^46^, nucleation-dependent scaffolding properties^40^, and protein conformational transitions^47^.

Our findings demonstrate that CRES has sub-micromolar affinity for DNA (Fig. 1). Like many other amyloid proteins which carry a net positive charge under physiological conditions^48^, the association is largely driven by electrostatic interactions between the basic polypeptide chain and acidic nucleic acid phosphate groups. Furthermore, previous studies of nucleic acid-binding amyloidogenic proteins have also observed binding affinities in the nanomolar-to-micromolar range resulting in elevated local protein concentration. This increased local concentration subsequently promotes protein aggregation through the formation of hydrophobic interactions between adjacent protein molecules^24,49,50^. Similarly, the stoichiometry of CRES-DNA complexes (Fig. 1) and associated increase in oligomerization kinetics (Fig. 4) reveal that multiple CRES molecules template off small DNA constructs similar to other amyloidogenic proteins (such as α-synuclein^51^) which have shown templating mechanisms of amyloid formation in physiological contexts. The rapid increase in oligomerization kinetics observed in the presence of DNA indicates that nucleic acid binding greatly facilitates CRES assembly (Fig. 4). In contrast to other previously studied amyloid-nucleic acid interactions, our work on DNA-mediated acceleration of CRES amyloid formation occurs in the context of two competing but complementary oligomerization pathways. These two paths could be a key factor in the regulated assembly of functional amyloids *in vivo,* potentially explaining how CRES maintains its non-pathological nature despite its amyloidogenic properties and highlights one of the many routes (i.e., templating off DNA) these proteins can take to form diverse biologically important structures with different morphologies and functions.

Indeed, this study further supports our previous work suggesting the existence of two distinct assembly pathways for CRES: the L1 loop-mediated pathway and the CRES loop pathway^11^. For wild type CRES, the rate of oligomerization is determined by a competition between the intermolecular CRES loop-CRES interactions and L1 loop-mediated oligomerization. The NMR data and mutational studies reveal that DNA binding occurs preferentially through the CRES loop and subsequent oligomerization proceeds predominantly via the L1 loop mechanism since the CRES loop becomes inaccessible due to DNA binding (Figs. 3,5). Therefore, DNA binding via the CRES loop diminishes the CRES loop pathway and accelerates the L1 pathway. The CRES loop driven oligomerization rate, which could lead to the increased proximity of CRES molecules, may be significantly reduced, providing evidence that, in the presence of small DNA constructs, the L1 loop is the predominant oligomerization pathway among the two possible mechanisms. Our results suggest that the formation of early oligomeric nuclei could take place via unique CRES loop interactions, but that slower large-scale oligomerization largely takes place via the L1 loop pathway. The ability of CRES to rapidly transition between monomeric and oligomeric states in response to DNA binding while maintaining controlled assembly is a remarkable feature of this functional amyloid. This dynamic equilibrium, coupled with the dual assembly pathways, may be key to CRES’s non-pathological nature and its diverse physiological roles in the epididymal environment.

The presence of extracellular nucleic acids in the epididymal amyloid matrix is necessary to stabilize matrix assembly and maintain the integrity of the amyloid infrastructure^13^. This stabilization is evidenced by the increased resistance to nuclease degradation observed in mature stages of the amyloid matrix in the distal part of the epididymis^22^. Nucleic acid-mediated CRES assembly could therefore enhance matrix structural organization and packaging, which may be crucial for its host defense functions and morphological plasticity. Additionally, the epididymal amyloid matrix may create a protective barrier that shields the germline from oxidative damage and other environmental stressors.

It is important to note that different sequences, size, and structure of nucleic acids might elicit outcomes distinct from what we observed here including differing rates of acceleration for CRES oligomerization and other amyloid morphologies. Indeed, extracellular DNA in other host defense amyloid structures such as bacterial biofilms ranges in size from small fragments to segments greater than 10 kb^52,53^. In addition, different eDNA structures including eDNA/eRNA hybrids have been found^54^. Further studies are needed to determine if and how different nucleic acid populations affect CRES oligomerization. Additional studies will also be important to determine in the presence of other environmental exposures such as bacterial virulence factors which assembly pathway predominates in CRES amyloidogensis including if the pathogenicity of the bacterial strain can result in the prioritization of one pathway over the over. For example, the presence of other external stimuli may drive amyloid assembly via the CRES loop allowing different morphologies to form (branched matrices instead of films and fibrils).

In summary, our detailed insights into DNA-mediated CRES oligomerization over a range of length and time scales highlight the importance of charge complementarity, increased local concentrations, and the competing but complementary amyloidogenic pathways in CRES assembly. Our biophysical and biochemical analyses supplement previous studies of the association of amyloidogenic proteins with nucleic acids and extend these results by providing an atomic-level, dynamic perspective for the oligomerization of functional amyloidogenic proteins such as CRES. The involvement of multiple pathways, rather than a single amyloidogenic core as observed in many pathological amyloids, likely plays a crucial role in conferring structural and functional heterogeneity to CRES. This represents nature’s elegant solution for generating diverse complexes and structures through multiple mechanistic pathways. These findings therefore offer significant potential for illuminating the complex mechanisms underlying biological amyloid assembly, their capacity to provide functional diversity in biological systems, and the profound influence of environmental signals on their resulting structures.

## MATERIALS AND METHODS

### Protein expression and purification

The pGEX-cs expression plasmid containing aa 20-142 and a C48A mutation of mouse CRES (Cst8) coupled to a TEV-cleavable N-terminal glutathione S-transferase (GST) protein^11^ was transformed into chemically competent *E. coli* Shuffle T7 cells (New England Biolabs). Transformed *E. coli* cells were grown at 30 °C in Luria broth (LB) or 2xM9 medium^55^ containing ^15^N ammonium chloride (1 g/L) and unlabeled D-glucose (3 g/L) as the sole nitrogen and carbon sources, respectively. The expression of GST-CRES was induced with 0.4 mM IPTG at OD_600_ ∼0.9-1.1 and growth continued at 12 °C for a maximum of 20 h in a shaking incubator. The cells were pelleted at 3750 x g, 4 °C for 30 min and resuspended in 40 mL lysis buffer (25 mM Tris, 250 mM NaCl, 1 mM EDTA, pH 8.0)/1 L cell pellet. Resuspended cells were incubated with 1 mM PMSF and 0.2 mg/mL DNase for 120 minutes before lysis by high pressure homogenization (Avestin C3). The soluble fraction containing GST-tagged CRES was collected by centrifugation at 39,000 x g at 4 °C for 45 min. The clarified lysate was loaded onto 20 mL/L culture of glutathione agarose resin (Pierce Glutathione Superflow Agarose, Thermo Scientific) equilibrated in lysis buffer and incubated overnight at 4 °C. The next day, the flow through was collected and the beads were washed with 25 mL of lysis buffer. GST-tagged CRES was eluted from the glutathione beads using 0.5% (w/v) reduced L-glutathione (Millipore-Sigma) in lysis buffer, adjusted to pH 7.4. The GST tag was removed from CRES by the addition of TEV protease at a ratio of 14 mg GST-tagged CRES to 1 mg TEV at 4 °C for at least 24 h. The NaCl concentration was then diluted to 41.6 mM using 25 mM MES, 1 mM EDTA, pH 6.0 before loading onto a HiTrap SP HP cation exchange column (Cytiva). A NaCl gradient was built from Buffer A (50 mM NaCl, 25 mM MES, 1 mM EDTA, pH 6.0) and Buffer B (1 M NaCl, 25 mM MES, 1 mM EDTA, pH 6.0) to separate CRES, GST, and TEV. Monomeric CRES was finally obtained through size exclusion chromatography on a HiLoad Superdex 200 pg column (Cytiva) in gel filtration buffer (25 mM MES, 250 mM NaCl, 1 mM EDTA, pH 6.0). Some mutants (LMD, K109/110E) were purified without the cation exchange step, because they did not bind to the HiTrap SP column, and were instead purified by serial HiLoad Superdex 200 size exclusion chromatography runs to separate the protein from the GST tag and TEV. The CRES mutants (Supplementary Table 1) were generated via site-directed mutagenesis using a modified Quikchange protocol (Millipore- Sigma) and were verified by Sanger sequencing.

For in vitro assays, approximately 3-4 mg of freshly S200-purified CRES in 25 mM MES, 250 mM NaCl, pH 6 were buffer exchanged into 25 mM MES buffer, pH 6 using an Amicon Ultra-15 10LJK centrifugal filter (Millipore). Monomeric CRES was isolated at 0.2 mg/mL; early oligomers were isolated by concentrating monomers to 2.0–2.5 mg/mL; and intermediate oligomers were formed by adjusting the early oligomers to pH 6.8 using 50 mM HEPES and further concentrating to 10–12.0 mg/mL using an Amicon Ultra-0.5 10 kDa MWCO centrifugal filter^13^. These different stages of assembly were then immediately used for subsequent experiments. All NMR samples were made with early oligomeric states (100-200 μM) of CRES in 25 mM MES pH 6.

### Fluorescence polarization assays for quantifying DNA binding

DNA binding affinities were measured by monitoring the change in fluorescence polarization (FP) of a fluorescently labeled 7-bp hairpin DNA (5’-/AF488/CACAAGCTTTTGCTTGTGAC-3’), 15-bp hairpin DNA (5’-/56- FAM/CACGCACGTAGAAGCTTTTGCTTCTACGTGCGTGAC-3’; IDT), or 40-bp dsDNA (5’-/6-FAM/GTGTTCGGACTCTGCCTCAAGACGGTAGTCAACGTGCTTG-3’) annealed to an unlabeled complementary strand; IDT). Hairpin DNA substrates were folded by denaturing at 95 °C for 5 min and snap cooling on ice. The 40-bp duplex was annealed by denaturing an equimolar ratio of the two strands at 95 °C for 5 min and then slowly cooling to room temperature. CRES was titrated into 21 µL reactions in 384-well black, flat bottom plates containing 100 nM of DNA substrate in buffer (25 mM MES pH 6.0 for monomer and early oligomer or 25 mM MES, 50 mM HEPES, pH 6.8 for intermediate oligomers). The plate was centrifuged for 1 min at 500 x g and was incubated for 10 min at room temperature. Post incubation, FAM FP was measured in a Biotek Synergy Neo2 plate reader using a FP 485/530 filter. FP data was fit to a simple two-state binding model and apparent K_d_s were calculated using the following equation:

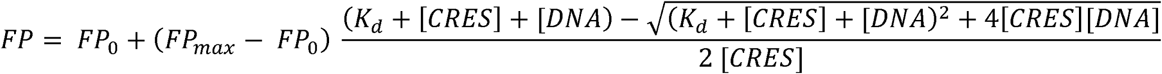

where [DNA] is the concentration of DNA, *K*_d_ is the dissociation constant, *FP_0_* is the FP in the absence of protein, and *FP_max_* is FP at maximum binding. Data points and error bars indicate the average and standard deviation of nLJ≥LJthree experiments on at least two different biological replicates per construct.

### Fluorescence polarization assays for quantifying stoichiometry

The stoichiometry of the CRES-DNA binding reaction was determined by titrating CRES into reactions containing excess unlabeled DNA (∼10 – 20 times the K_d_) and 100 nM of the fluorescently labeled DNA constructs described above. Specifically, increasing concentrations of purified CRES were titrated into 21 µL reactions in 384-well black, flat bottom plates containing 5 µM of either hairpin or dsDNA substrates and 25 mM MES pH 6.0 (monomeric conditions). The plate was centrifuged for 1 min at 500 x g and was incubated for 10 min at room temperature. Post incubation, FAM FP was measured in a Biotek Synergy Neo2 plate reader using the FP 485/530 filter. The data was analyzed by finding the intersection point between the linear increase in FP and saturation in FP, which corresponds to the ratio of protein:DNA where all of the DNA is bound.

### Dynamic light scattering

Monomeric CRES samples ranging from 0.17-0.22 mg/mL in 25 mM MES, pH 6.0 were examined by DLS. 350-400 μL of monomeric CRES with or without equimolar 15-bp hairpin DNA were placed in a UV-Cuvette (ZEN0118, Malvern) and analyzed at 25 °C using a Malvern Zetasizer (Model: ZMV2000). “Size” and “Protein” were selected as the measurement and the material types, respectively. For each sample, the measurement was repeated at least 3 times. Each time, 15 scans were acquired, and each scan lasted for 15 s. After the diffusion of a particle moving under Brownian motion was measured, the Zetasizer software version 7.13 converted the diffusion to a size and generated size-intensity distributions using Stokes-Einstein relationship with a refractive index of 1.33 nD and dynamic viscosity of 0.8872 cP. Gaussian fitting was then used to extract the size mean. The analysis was done using the 300 size classes with the lower and upper size limits set to 0.4 nm and 10000 nm, respectively.

### Congo red assay

Congo red (Millipore-Sigma) was dissolved in 4 mM potassium phosphate pH 7.4 to make a 630 μg/mL stock. In a 384 well plate, a 20 μL reaction mixture was made with buffer (25 mM MES pH 6.0 or 25 mM MES, 50 mM HEPES, pH 6.8) and CRES (monomer/early oligomer/intermediate oligomer) with or without equimolar 15-bp hairpin DNA. 1 μL of Congo red stock was added to the reaction mixture, making the final concentration of Congo red in each well 30 μg/mL. The plate was centrifuged for 1 min at 500 x g and was incubated for 10 min at room temperature. Fluorescence measurements were performed in a Biotek Synergy Neo2 plate reader with excitation at 525 nm and emission at 625 nm. 10 kinetic reads were measured every 30 s with shaking for 10 s between every read. The relative fluorescence units were calculated by subtracting buffer controls within each read and averaging the 10 reads. Data points and error bars indicate the average and standard deviation of nLJ≥LJthree experiments on at least two different biological replicates per construct.

### Transmission electron microscopy

A 900 µM stock of 15-bp hairpin DNA in 25 mM MES, 25 mM NaCl, pH 6 was heated at 95 °C for 2 min to remove secondary structure, immediately placed on ice, and then aliquoted and stored at -25 °C until use. 9.4 µM CRES was incubated with equimolar concentrations of 15-bp hairpin DNA in 25 mM MES pH 6 at room temperature for up to 2 days. Control samples included 9.4 µM CRES alone and 9.4 µM 15-bp hairpin DNA alone in 25 mM MES pH 6. After 60 min incubation (day 0), 24 hrs (day 1) and 48 hrs (day 2), 5 µL was removed from each sample and spotted on to formvar/carbon coated 200 mesh nickel grids (Ted Pella, Redding, CA) for 5 min. The sample was wicked off with filter paper, washed for 1 min with water, stained with 2% uranyl acetate for 1 min, and washed again with water for 1 min. Samples were allowed to dry for 1 min. Images were captured using a Hitachi transmission electron microscope.

### NMR experiments

NMR data was collected at 25 °C using Bruker 850 MHz Avance III (19.97 T) and Bruker 600 MHz Avance NEO (14.1 T) spectrometers equipped with 5 mm TCI cryoprobe with z-axis gradient. NMR data were processed with NMRPipe/NMRDraw^56^ and analyzed with CCPN analysis v2.5^57^. Backbone amide ^1^H and ^15^N assignments for wild type CRES were taken from previous studies^11^. DNA titration experiments were performed by adding 7-bp or 15-bp hairpin DNA to ^15^N-labeled CRES and monitoring the change in amide chemical shifts in 2D ^15^N,^1^H TROSY-HSQC^58^ spectra until complete saturation was reached. Amide chemical shift perturbations (CSPs) were calculated as:

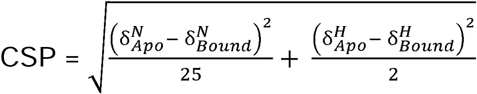

where δ*^N^* and δ*^H^* are the nitrogen and proton chemical shifts, respectively.

Backbone ^15^N R_1_ and R_1ρ_ for wild type and mutant CRES in the absence and presence of 2.5 or 5% mol/mol 15-bp hairpin DNA were also acquired at 600 MHz and 850 MHz at 25 °C (300 µM CRES wild-type, 460 µM CRES T79G, 380 µM CRES T79G+LMA). ^15^N R_1_ and R_1ρ_ relaxation rates were calculated from the single exponential fits of the intensities from six and seven parametrically varied time points ranging from 2–200 msec (R_1_) and 2–60 msec (R_1ρ_), respectively. R_1ρ_ values were converted to R_2_ using the relation:

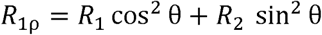

where 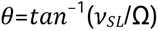 is the effective tilt angle of the rotating frame, *v_sl_* is the field strength of the applied spin lock (2000 Hz), and Ω is the offset of the peak from the ^15^N carrier. Errors in the relaxation rates were calculated from the covariance matrix of the fit.

When a small, fast tumbling molecule (e.g., CRES) is exchanging with a much larger, slowly tumbling molecule (e.g., CRES_n_-DNA), the exchange kinetics (i.e., the exchange rate – k_ex_) and thermodynamics (i.e., population of the lowly populated larger complex – p_b_) can be established from dependence of the ^15^N ΔR_2_ (R_2,+DNA_ – R_2,apo_) on the strength of the static magnetic field B_0_.

A quantitative measure of field dependence of ΔR_2_ at 600 and 850 MHz is given by^28,32^

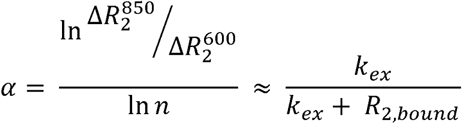

where n is the theoretical factor by which ΔR_2_ is expected to increase between data collected at 600 MHz and 850 MHz field strengths (n = 1.25) and R_2,bound_ is the transverse relaxation rate for the slowly tumbling CRES_n_-DNA state. When exchange is fast on the timescale of transverse relaxation (i.e., 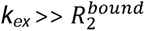),

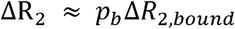

Comparisons between the ^15^N R_2_ and ^15^N ΔR_2_ for different samples were made and analyzed through GraphPad Prism. Statistical analysis was done using one-way ANOVA involving comparison between the means of multiple unmatched groups with the pooled standard deviations of the groups. Tukey’s multiple comparisons test was performed as a post-test to compare all possible pairs of data controlling the family-wise error rate.

#### HADDOCK

Molecular docking of the 15-bp hairpin DNA to monomeric CRES was performed with the GURU interface of the HADDOCK v2.4 webserver^26,27^. For the two-body rigid-body docking protocol, the PDB inputs were the crystal structure of monomeric CRES (6UIO)^11^ and a 15-bp hairpin DNA modeled through MD simulations^59^. CRES residues which showed significant chemical shift perturbations in intermediate exchange in the NMR titration experiments were used as ‘active’ ambiguous restraints (residue numbers 10, 15, 18, 25, 26, 33, 34, 36, 39, 43, 49, 54, 58, 61, 68, 69, 70, 71, 73, 76, 89, 91, 92, 93, 95, 97, 99, 101, 104, 105, 113, 117), neighboring CRES residues were automatically chosen as ‘passive’ ambiguous restraints by HADDOCK, all the DNA residues were set to ‘active’ ambiguous restraints. 50% of the ‘active’ and ‘passive’ restraints were omitted for each run, and all default settings were used otherwise.

For docking the domain-swapped dimer model for CRES, two N-truncated models of CRES monomer were cleaved at the L1 hinge, generating four polypeptides in the docking calculation. Active residues were defined along the antiparallel β-sheet interactions in the β2-L1-β3 region between the two monomeric units (Monomer1_14to79: 67, 68, 69, 70, 71, 72, 73, 74, 75, 76, 77, 78. (Partner Selection 4). Monomer1_80to142: 81, 82, 83, 84, 85, 86, 87, 88, 89, 90, 91, 92, 93 (Partner Selection 3). Monomer2_14to79: 67, 68, 69, 70, 71, 72, 73, 74, 75, 76,77,78 (Partner selection 2). Monomer2_80to142: 81, 82, 83, 84, 85, 86, 87, 88, 89, 90, 91, 92, 93 (Partner selection 1). Unambiguous restraints (derived from the crystal structure of wild-type CRES PDB: 6UIO) included interactions between α-helix and β-sheets of the two domains and the antiparallel β-sheet interactions of the two L1 hinges that convert to β-strands (Supplementary Table 2). Hydrogen bond restraints were used to join the four constructs together according to the sequence of the two monomeric domains (Supplementary Table 3). C2 symmetry was also enforced between the two CRES monomers to allow HADDOCK to move the two monomers relative to each other. This model was then used further in a two-body docking calculate to model a domain-swapped dimer DNA-bound complex. The same active residues were used as used above for the monomeric CRES-DNA docked models. PyMOL v3.0 was used to make figures of structures.

## Data availability

Data are available from the corresponding author upon reasonable request.

## Acknowledgements

We are grateful to M. Canny for critically reviewing this manuscript. This work was supported by NIH/NIGMS (R35GM128906 to M.P.L.) and NIH/NIA (R21AG089761 to G.A.C.).

## Author Contributions

R.K., G.A.C., and M.P.L. designed the research. R.K. prepared protein samples and conducted DNA binding and NMR experiments. G.A.C. prepared protein samples and conducted TEM experiments. R.K., G.A.C., and M.P.L. analyzed the data. R.K. drafted the paper. R.K., G.A.C., and M.P.L. edited and approved the final manuscript.

## Notes

### Competing Interest Statement

The authors have declared no competing interest.

