## Supplemental Information for "DNA modulates structural transitions and oligomerization kinetics of the functional amyloid CRES"

**Supplementary Tables:**

**Supplementary Table 1: CRES mutants and their effects on DNA binding**

| Mutant | Description | Effects on DNA binding |
|---|---|---|
| **LMD:** N94D/N101D/T102D/N104D | CRES loop charge reversal mutant | DNA binding decreased |
| **K109E/K110E** | CRES loop charge reversal mutant | DNA binding decreased |
| **K75E** | L1 loop charge reversal mutant | DNA binding decreased |
| **R81E** | L1 loop charge reversal mutant | DNA binding decreased |
| **LMA:** N94A/N101A/T102A/N104A | CRES loop pathway blocking mutant | No effect on DNA binding. Binds like wild type |
| **T79G** | L1 loop pathway blocking mutant | No effect on DNA binding. Binds like wild type |
| **T79G+LMA:** T79G/N94A/N101A/T102A/N104A | CRES loop and L1 loop pathway blocking mutant | No effect on DNA binding. Binds like wild type |

**Supplementary Table 2: Unambiguous restraints for domain-swapped dimer modelling in HADDOCK**

| Chain A/C Residues | Chain B/D Residues | Distance (Å) |
|---|---|---|
| 79 | 80 | 1.3 |
| 40 | 71 | 3.9 |
| 40 | 74 | 3.9 |
| 45 | 74 | 3.9 |
| 45 | 71 | 3.9 |
| 49 | 71 | 3.9 |
| 46 | 71 | 5 |
| 44 | 84 | 3.1 |
| 48 | 86 | 3.9 |
| 56 | 90 | 3.7 |
| 49 | 88 | 3.7 |
| 57 | 66 | 1.8 |
| 60 | 66 | 3.7 |
| 56 | 121 | 3.6 |
| 52 | 121 | 3.8 |
| 51 | 123 | 5.2 |
| 56 | 119 | 3.2 |
| 56 | 117 | 5.1 |
| 66 | 117 | 4.4 |
| 47 | 132 | 8.1 |
| 66 | 115 | 5.7 |
| 51 | 132 | 4.1 |
| 51 | 134 | 3.8 |
| 59 | 137 | 1.8 |
| 56 | 137 | 3.7 |
| 56 | 139 | 3.5 |
| 48 | 132 | 4.1 |

**Supplementary Table 3: Hydrogen bond restraints for domain-swapped dimer modelling in HADDOCK**

| Chain A Residues | Chain B Residues | Chain C Residues | Chain D Residues | Distance (Å) |
|---|---|---|---|---|
| 79 | 80 | | | 1.3 |
| | | 79 | 80 | 1.3 |
| 57-66 | | | | 1.3 |
| | | 57-66 | | 1.3 |
| 59 | | | 137 | 1.3 |
| | 137 | 59 | | 1.3 |

### Supplementary Figures:

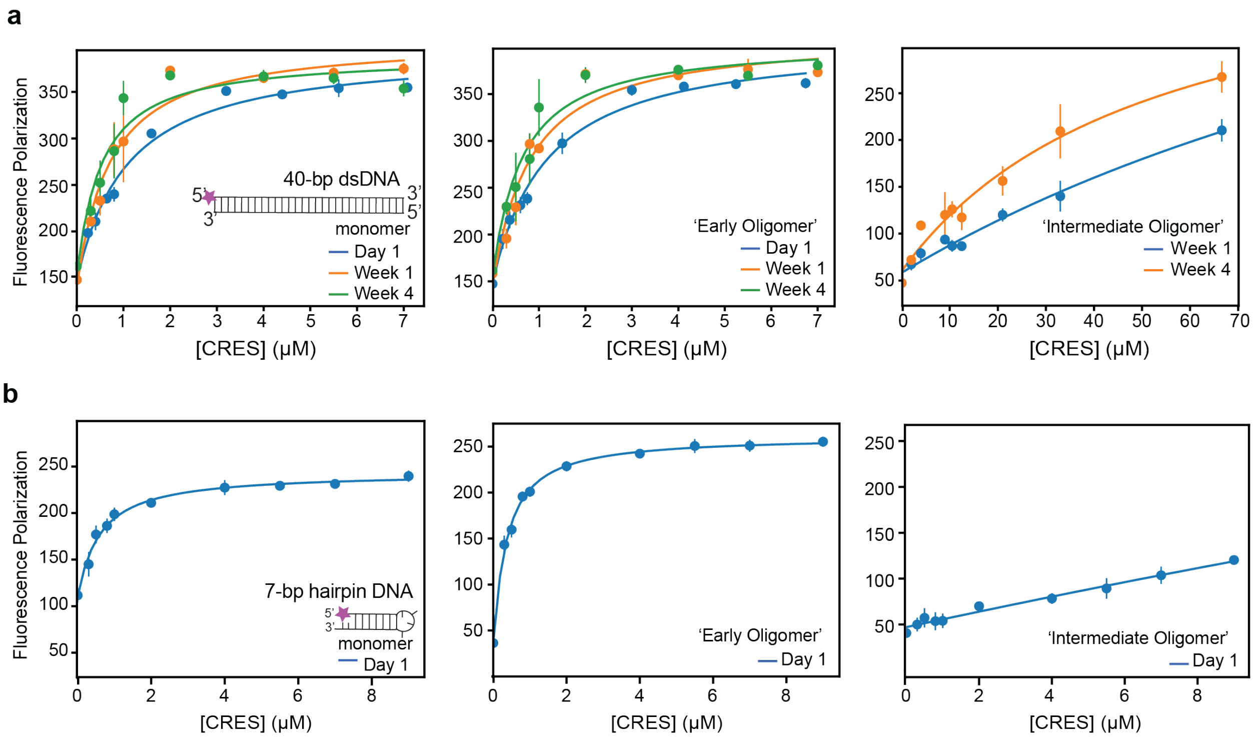


**Supplementary Figure 1. CRES binds to DNA with nanomolar affinity.** Change in polarization of a fluorescently labeled (a) 40-bp dsDNA or (b) 7-bp hairpin DNA as a function of the concentration of CRES for different stages of assembly. The data points are displayed as the mean and standard deviation of three experiments, and the solid lines represent the fitting of the data to a two-state binding isotherm.

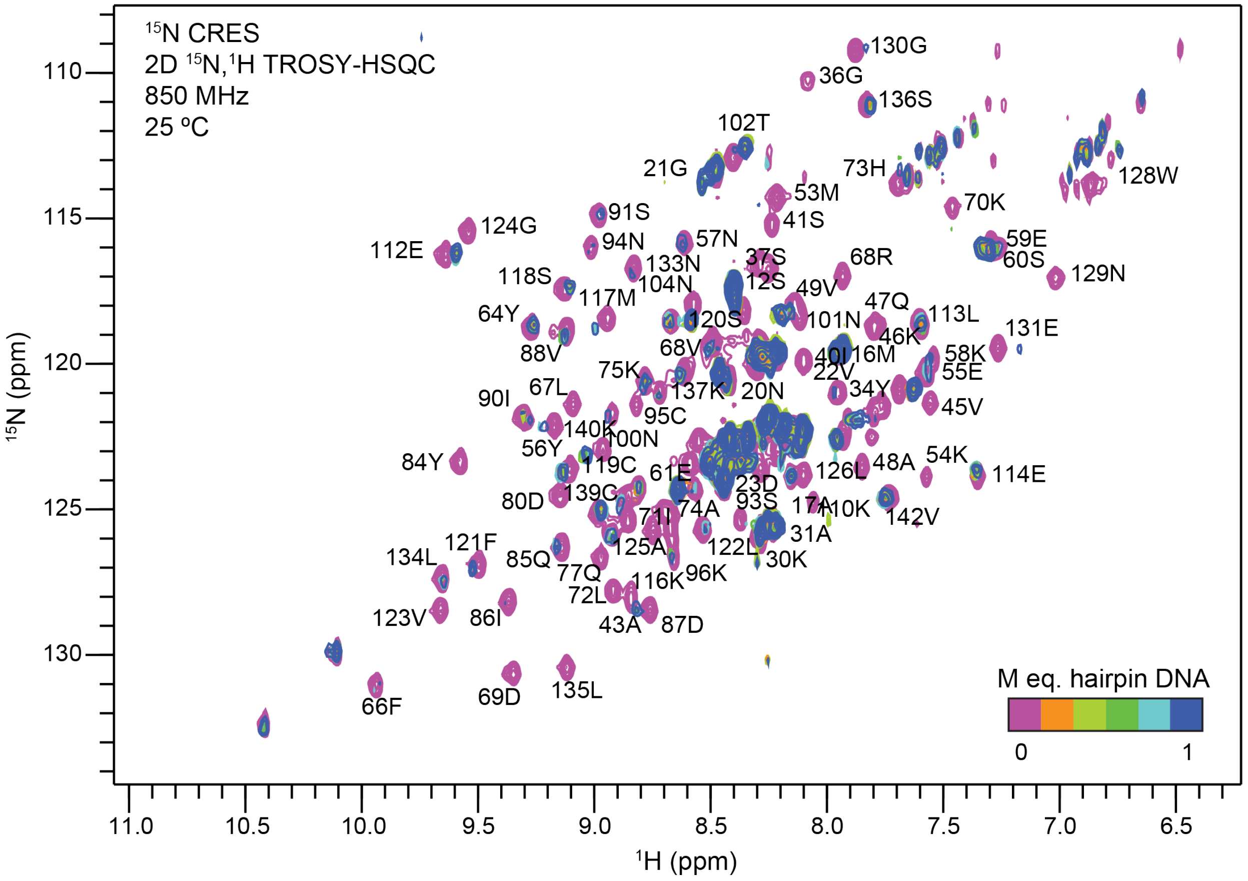


**Supplementary Figure 2. NMR titration of CRES with a 15-bp hairpin DNA.** Overlay of the 2D $^{15}$N,$^{1}$H TROSY-HSQC spectra of $^{15}$N-labeled CRES with increasing concentrations of 15-bp hairpin DNA.

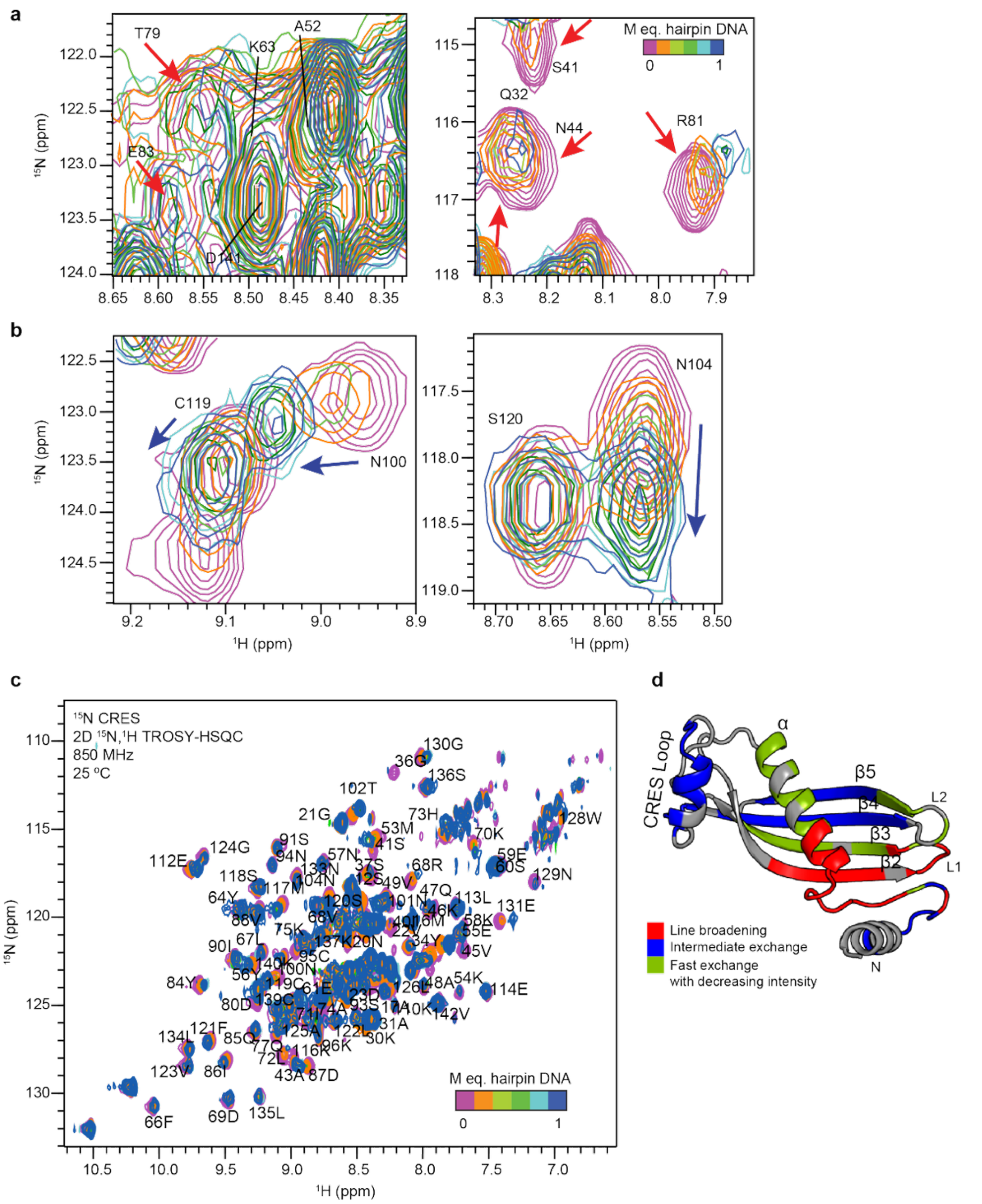


**Supplementary Figure 3. NMR titration of CRES with a 7-bp hairpin DNA.** (a and b) Overlay of regions from the 2D $^{15}$N,$^{1}$H TROSY-HSQC spectra of $^{15}$N-labeled CRES with increasing concentrations of 7-bp hairpin DNA. The overlaid spectra show resonances undergoing moderate line-broadening (a; highlighted by red arrows), and resonances undergoing intermediate exchange (b; highlighted by blue arrows). (c) Overlay of the entire 2D $^{15}$N,$^{1}$H TROSY-HSQC spectra of $^{15}$N-labeled CRES with increasing concentrations 7-bp hairpin DNA. (d) Perturbations mapped onto the monomeric structure of CRES (PDB ID:6UIO)[11] illustrating residues that experience line-broadening (red), intermediate exchange (blue), or fast exchange (green).

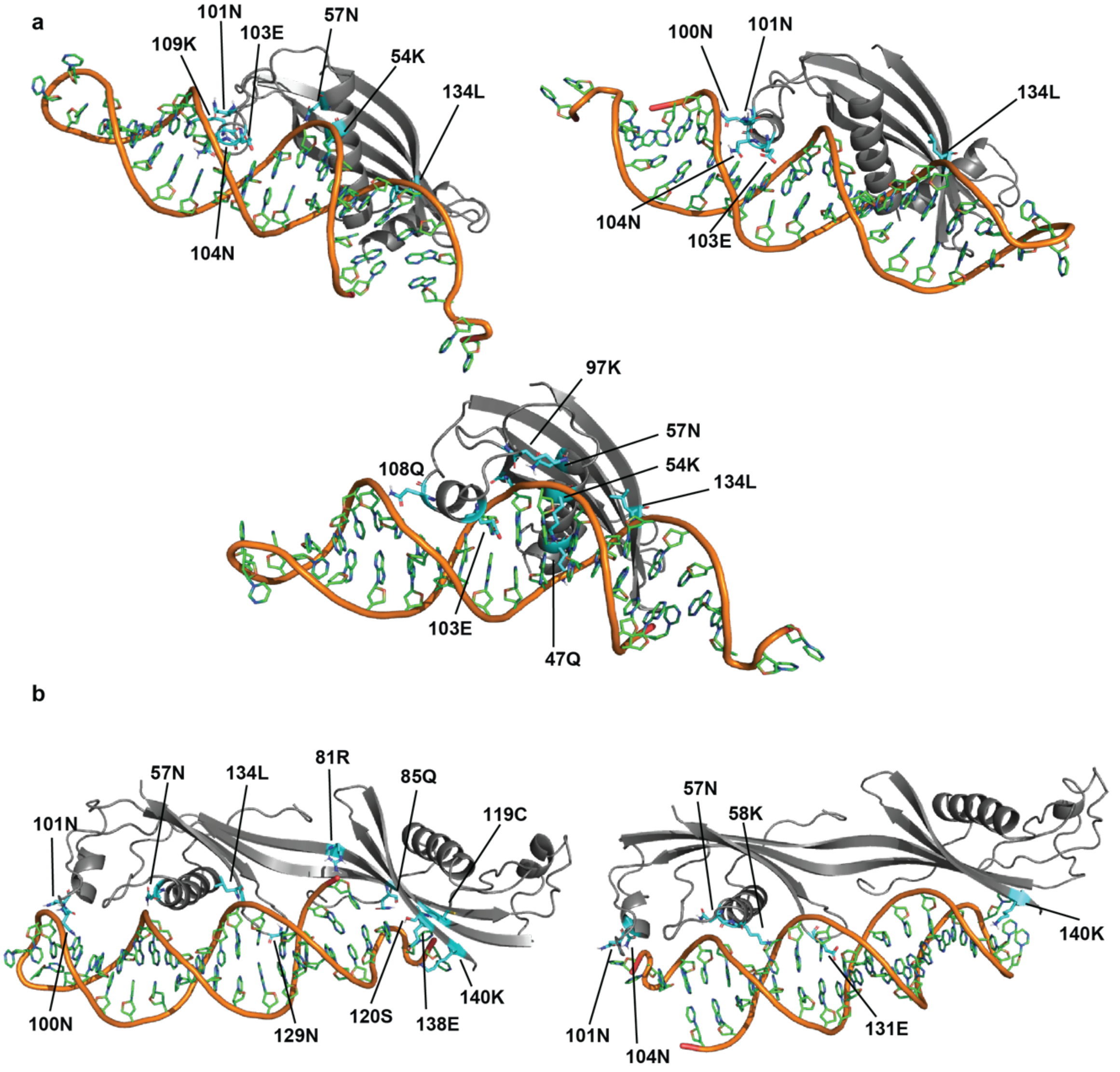


**Supplementary Figure 4. Preliminary models of DNA bound CRES.** HADDOCK 2.4 generated models of (a) monomeric CRES bound to 15-bp hairpin DNA and (b) L1/domain-swapped dimer CRES bound to 15-bp hairpin DNA. Residues in cyan sticks highlight polar interactions between the protein and DNA.

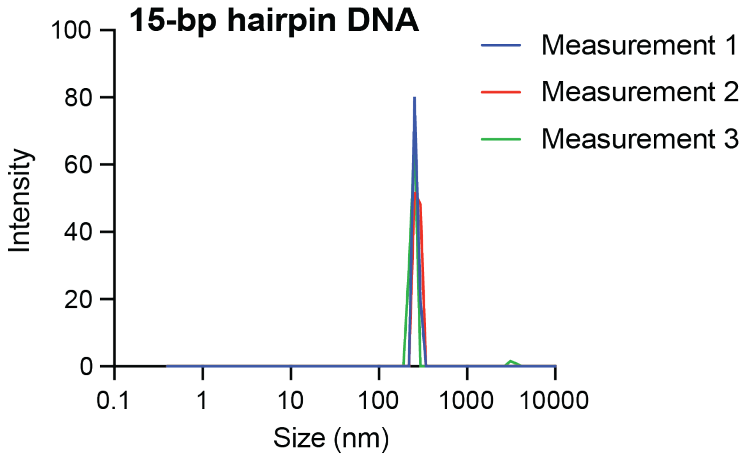


**Supplementary Figure 5. Dynamic light scattering data for the 15-bp hairpin DNA only control.**

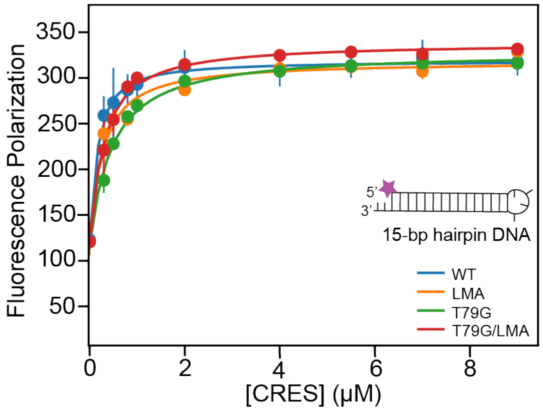


**Supplementary Figure 6. CRES pathway blocking mutants do not affect DNA binding.** (a) Change in polarization of a fluorescently labeled 15-bp hairpin DNA as a function of the concentration of monomeric CRES mutants. The data points are displayed as the mean and standard deviation of three experiments, and the solid lines represent the fitting of the data to a two-state binding isotherm.

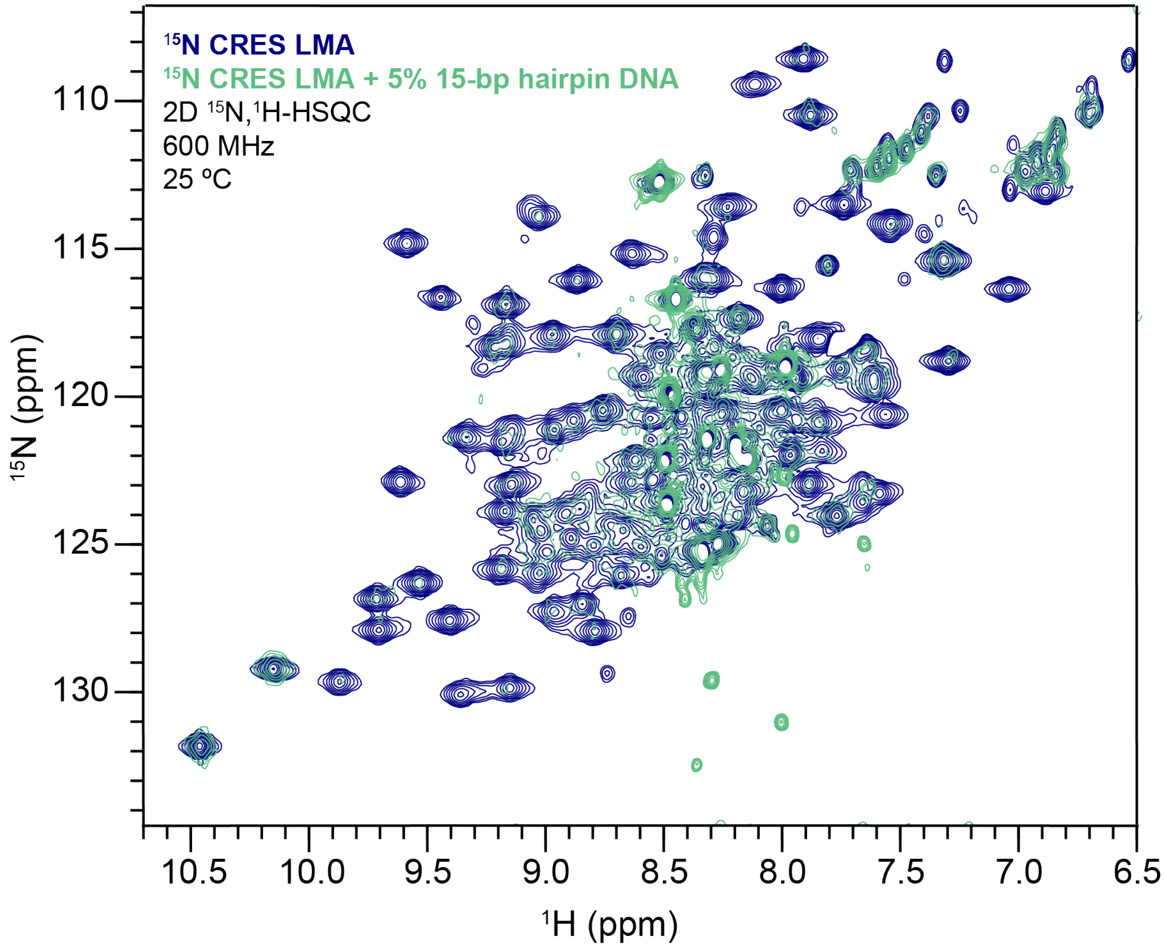


**Supplementary Figure 7. CRES LMA mutant exhibits extreme line-broadening in the presence of sub-stoichiometric amounts of DNA.** (a) Overlay of 2D $^{15}$N,$^{1}$H TROSY-HSQC spectra of $^{15}$N-labeled CRES LMA in the absence (blue) and presence (green) of 5% 15-bp hairpin DNA.

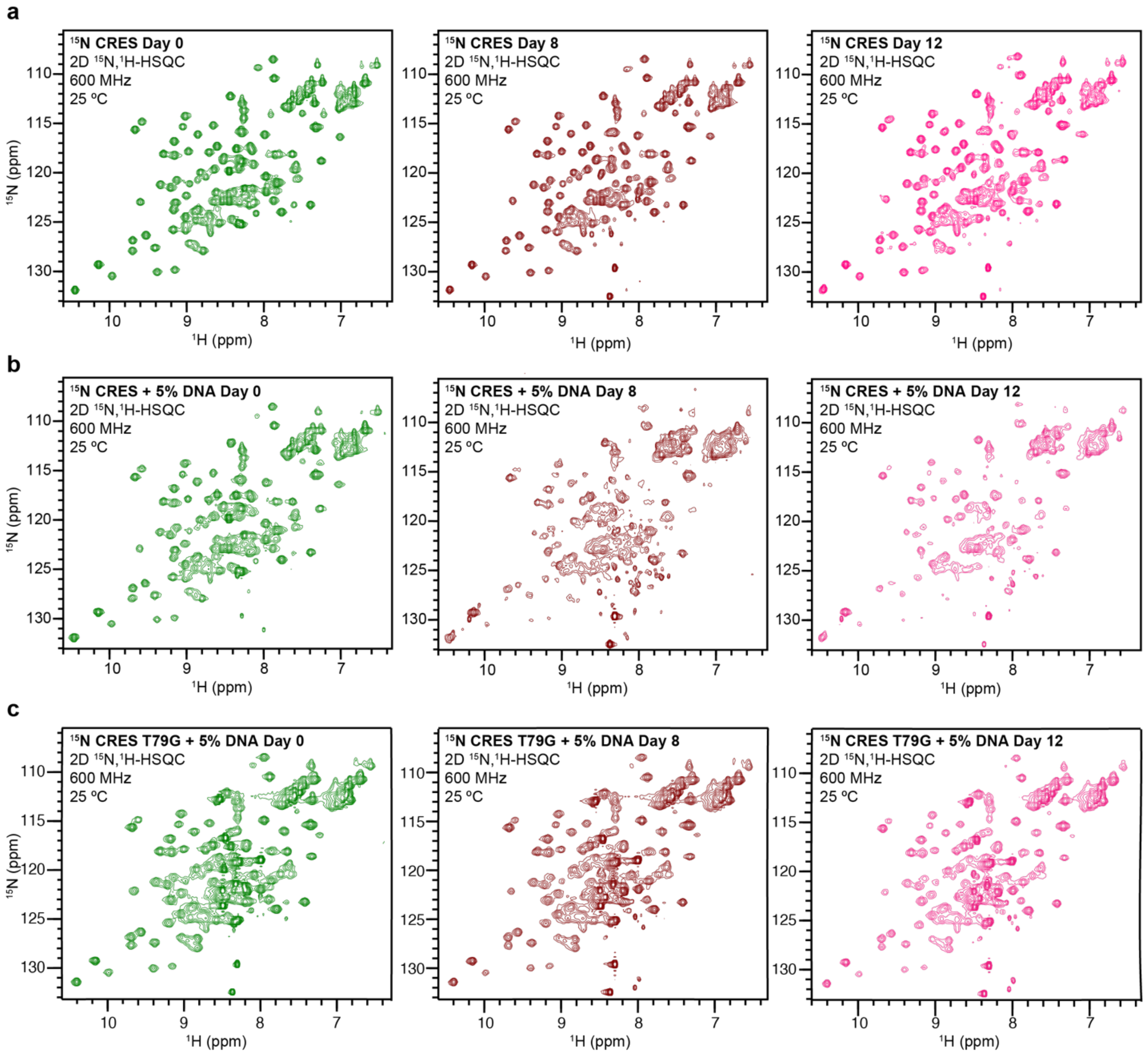


**Supplementary Figure 8. CRES shows rapid NMR intensity loss in the presence of DNA.** 2D $^{15}$N,$^{1}$H HSQC spectra of $^{15}$N-labeled CRES in the absence of DNA (a) or with 5% 15-bp hairpin DNA (b), and $^{15}$N-labeled CRES T79G incubated with 5% 15-bp hairpin DNA (c) at 0, 8, and 12 days (green, brown, and pink, respectively).
